# Metabolic state modulates risky foraging behavior via multiple branches of the insulin/IGF-1-like pathway in *C. elegans*

**DOI:** 10.1101/2025.02.24.639891

**Authors:** Kaiden H. Price, Jason T. Braco, Preeti F. Sareen, Peter J. Niesman, Michael N. Nitabach

## Abstract

Foraging to acquire nutrients is an essential and sometimes risky behavior displayed by nearly all animals. Appropriately balancing foraging risks with nutrient requirements is pivotal for peak survival and reproduction, and metabolic state (i.e., how urgently the animal requires nutrients) is a strong modulator of risky foraging behavior. In this study, we asked what molecular signal allows *C. elegans* to change its foraging behavior in response to changes in its metabolic state. We used an assay of risky foraging behavior, where wild type worms increase risky foraging behavior after food deprivation, to screen for candidate genes. We found that DAF-2, the singular receptor in the *C. elegans* insulin/IGF-1 signaling (IIS) pathway, is necessary for worms to modulate risky foraging behavior in response to short-term food deprivation. Worms with mutations in genes upstream and downstream of *daf-2* in the IIS pathway also exhibited a reduction in the effect of food deprivation. While a canonical understanding of the IIS pathway would suggest that the FOXO transcription factor DAF-16 is the primary downstream IIS pathway target, we found that DAF-16 was not required for worms to exhibit food-deprivation-driven changes in foraging behavior. Furthermore, we determined that the calsyntenin ortholog CASY-1, which allows DAF-2c to traffic to axons, is required for food deprivation to modulate risky foraging behavior. These results both validate the IIS receptor as a pivotal regulator of risky foraging behavior and suggest a multi-pronged downstream pathway. Overall, these data enrich our understanding of how organisms transduce metabolic state information to make vital decisions about when to engage in risky foraging behaviors.

**ARTICLE SUMMARY:** *C. elegans* changes its foraging behavior from a low-risk to a high-risk strategy when it is food deprived. Our paper demonstrates that the DAF-2 insulin/IGF-1-like receptor is an essential modulator for this behavior following food deprivation. Moreover, the regulation of risky foraging behavior is enacted by multiple downstream effectors of the insulin/IGF-1-like pathway, including a canonical pathway involving the FOXO transcription factor DAF-16 and a non-canonical pathway involving CASY-1, which can traffic an isoform of DAF-2 to axons.

## INTRODUCTION

Almost all animals forage – they display distinct behavioral patterns to search for and acquire nutrients (Kuburich et al. 2016; Rudebeck and Izquierdo 2022; Zjacic and Scholz 2022). However, even for the simplest of organisms, foraging is a complex trade-off between risk and reward. Typically, conservative foraging strategies (e.g. staying in one place) are less likely to result in nutrient acquisition; however, more risky foraging strategies cost energy and increase the likelihood that the animal will encounter environmental hazards, predation, and injury. Effectively balancing nutrient requirements with foraging risks requires ongoing assessment of the external environment (nutrient sources, predation risks, environmental hazards, etc) and the internal environment (metabolic state, nutrient deficiencies, developmental stage, etc). The animal must then integrate that information to drive appropriate behavioral choices (Lima and Dill 1990; Mukherjee and Heithaus 2013; Zjacic and Scholz 2022).

In humans, foraging behavior now occurs in the context of a broad array of human cultures and agricultural systems (Capocasa and Venier 2024), and human behaviors around seeking and obtaining food are commonly understood as dietary choice behaviors (Wadi et al. 2024). These dietary choices impact a wide variety of health outcomes, including body fat composition, cancer risk, heart health, and risk of dementia (Guasch-Ferré and Willett 2021; Clemente-Suárez et al. 2023). Body mass index, or BMI (a flawed but frequently-used measure that estimates body fat on the population level (Gonzalez et al. 2017; Bray 2023)) has been found to be 30-55% heritable (Bouchard 2021), suggesting that internal physiologic drivers may deeply influence human feeding behaviors. Additionally, surgical and pharmacological interventions have shown substantially greater effects on weight loss than behavioral interventions alone (Blüher et al. 2023), further suggesting that a significant part of human foraging behavior is driven by physiological signals (Rebello and Greenway 2016; Watts et al. 2022). Expanding our understanding of how foraging behavior is regulated at the molecular level may lead to better interventions that can alter human dietary choices in ways that improve human health.

We chose to study the regulation of foraging behavior in the microscopic nematode worm *C. elegans.* This worm is an excellent model system for investigating the neural, genetic, and molecular mechanisms of behavior. Substantial work has already been done to examine the regulation of feeding behavior in *C. elegans*, identifying relevant neurons (Gray et al. 2005; Ghosh et al. 2016; Rhoades et al. 2018), neurotransmitters (Flavell et al. 2013; Lee et al. 2017; Gadenne et al. 2022), and genes (Matty et al. 2022; McLachlan et al. 2022; Boor et al. 2024). However, the molecular mechanism by which *C. elegans* integrates metabolic state into foraging decisions is not yet fully understood.

We previously employed a multisensory decision assay that is well-suited for assessing foraging risk tolerance in *C. elegans* (Ghosh et al. 2016). This assay presents worms with a complex choice: cross a hyperosmolar barrier to approach a food odor, or avoid the hyperosmolar barrier and thus fail to approach the food odor (Ishihara et al. 2002; Shinkai et al. 2011). While well-fed worms overwhelmingly avoid both the hyperosmolar barrier and the food odor, worms that have been food deprived will generally cross the hyperosmolar barrier and move towards the food odor (Ishihara et al. 2002; Ghosh et al. 2016; Matty et al. 2022). Here, we used this foraging risk assay to examine which genes may be involved in allowing the worm to integrate its internal metabolic state (i.e. whether it is well-fed or food deprived) into its foraging behavior. We initially ran a mutant screen for candidate genes using established mutant worm lines. This candidate screen had two clear hits: *egl-3* and *daf-2*. We chose to focus on *daf-2*, the insulin/IGF-1-like receptor in *C. elegans*.

The canonical insulin/IGF-1 signaling (IIS) pathway in *C. elegans* first activates or antagonizes the DAF-2 receptor with one or more of forty ILPs (insulin-like peptides) (Zheng et al. 2018). The cell then transmits the signal from the DAF-2 receptor via a series of kinases, beginning with AGE-1, the ortholog of mammalian PI3K (Weinkove et al. 2006). AGE-1 phosphorylates PIP2 to increase concentrations of PIP3, which activates downstream kinases; DAF-18, a PTEN ortholog, opposes this activity, dephosphorylating PIP3 and decreasing activating of downstream kinases (Murphy 2013). The insulin receptor substrate protein, IST-1, facilitates AGE-1 activity, and may act to pair non-canonical effectors to DAF-2 receptor activity (Murphy 2013). Ultimately, the AKT kinases phosphorylate the transcription factor DAF-16 (orthologous to the mammalian FOXO) (Paradis and Ruvkun 1998). When phosphorylated, DAF-16 is sequestered in the cell soma, where it is unable to act as a transcription factor (Murphy 2013). We found that many of the genes in the IIS pathway in *C. elegans* are necessary for the worm to adjust its foraging behavior in a way that appropriately discriminates between a well-fed and a food-deprived metabolic state. Moreover, while previous research supports a role for *daf-2* in modulating foraging behavior (Matty et al. 2022), our research further suggests that two downstream IIS pathway elements (the FOXO transcription factor DAF-16 and the calsyntenin ortholog CASY-1) are likely acting in two parallel pathways to modulate foraging risk tolerance.

## MATERIALS AND METHODS

### Strains and Culturing

Worms were maintained at 20C on NGM media cultured with *E. coli* OP50 as a food source. All strains were obtained from the Caenorhabditis Genetics Center. The Bristol N2 strain was used as wildtype (WT) in all assays. In all assays, worms were age-matched by picking L4 worms 18-24 hours before assays. As many *daf-2* strains form dauers at 20C (Gems et al. 1998), all *daf-2* strains were maintained at 15C alongside paired WT control plates. For these strains, L4 worms were picked and moved to 20C 18-24 hours before being used in assays. (In experiments where the mutant worms were maintained at 15C, the respective N2 WT control worms were also maintained at 15C).

### Foraging risk assay and osmotic avoidance assays

Multisensory foraging risk assays were conducted as in Ghosh *et al*. 2016. Plates were first prepared by taking fresh (no more than 5 day old) 60 mm NGM plates and drying the open plates in a closed chemical hood for 1 hour. A 1 cm diameter ring was drawn in the middle of the dried NGM plate using 10 µl 3M fructose (shown in pink in **Fig. 1a**). Two 1 µl dots of 1:350 diacetyl were then placed on opposite sides of the plate, 1 cm from the ring border (shown in purple in **Fig. 1a**). The plate was then set to the side for five minutes to allow an odor gradient to form. Meanwhile, 10 worms were moved from the culture plate to a clean NGM plate with no *E. coli* and left for at least 1 minute to allow any remaining bacteria to wash off onto the clean plate. At the end of the 5 minutes, the 10 worms were moved from the cleaning plate to the center of the assay plate inside the fructose ring. The worms were allowed to forage for 15 minutes, and then the number of worms remaining in the ring were counted. “Percent exiting” was determined as (10 – (# worms remaining)) * 10. (Unisensory osmotic avoidance assays were performed in essentially the same manner as our foraging risk assay, except that the diacetyl was not added to the assay plate.)

**Figure 1:**
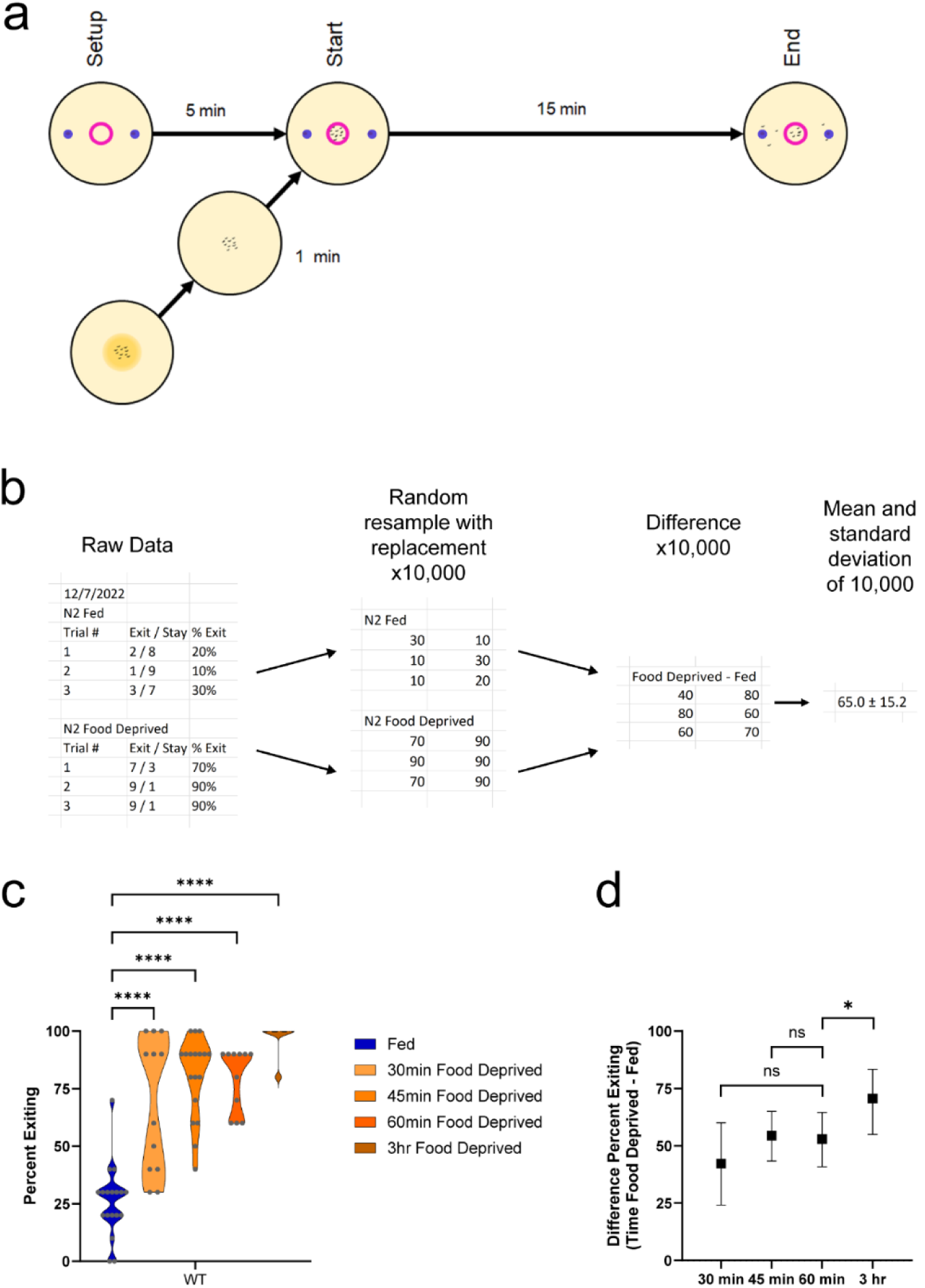
Assay and experimental design. a) A 1cm diameter circle of 3M fructose (represented by the pink circle) was drawn in the center of a 60 mm Petri dish of nematode growth media (NGM). Two dots of 1:350 diacetyl were placed outside the fructose circle, 1 cm from the edge of the Petri dish (represented by the purple dots). The dish sat for 5 minutes to allow an odor gradient to form. Meanwhile, 10 worms were moved from a feeding dish to a clean NGM dish for one minute to wash off any bacteria adhering to the worms. At the end of the 5-minute set up, the 10 worms were moved to the center of the fructose circle. Worms were allowed to forage for 15 minutes, and then the number of worms remaining in the circle was counted. b) Raw data was additionally analyzed by calculating the bootstrapped distribution of the difference between the fed and food deprived groups. Raw datasets of fed and food deprived worms were first transformed into bootstrapped datasets with 10,000 replications using resampling with replacement (the first two of 10,000 replications are shown here). A third bootstrapped dataset was created by calculating the difference of each data point from the fed and food deprived bootstrapped datasets. This third bootstrapped dataset of the difference (food deprived minus fed) was used to calculate a population average, standard deviation, and 95% confidence interval for the effect of food deprivation. c) Foraging behavior of fed and food deprived wild type worms in the foraging risk assay with varying durations of food deprivation (analyzed by one-way ANOVA with Dunnett’s post hoc tests, using ‘Fed’ as the control condition). d) Bootstrapped distributions (means and 95% confidence intervals) of the differences between fed and food deprived wild type worms from 1c (calculated as described in 1b; analyzed by one-way ANOVA with Dunnett’s post hoc tests, using ‘60 min Food Deprived difference’ as the control condition). Adjusted p-values of post hoc tests: ns, p >= 0.05; *, p < 0.05; ****, p < 0.0001.

### Food deprivation

For all assays that used food deprivation, worms were moved from a culture plate to a cleaning plate (as above) for at least one minute, and then placed into a food deprivation plate. The food deprivation plate was a clean, dried NGM plate with an approximately 4 cm diameter ring-shaped acetate stencil pressed gently into the agar to form a seal. The stencil reduced the number of worms that crawl up the side of the petri dish and desiccate. Worms remained in the food deprivation plate for 1 hour (unless otherwise specified), and then were moved directly onto the assay plate.

### Unisensory chemotaxis assay

Chemotaxis assays were performed as in Ghosh *et al*. 2016. Fresh 60 mm NGM plates were prepared with 1ul of 1:10,000 diacetyl in dH2O placed 1 cm from one edge of the plate, and 1ul of dH2O placed 1 cm from the opposite edge of the plate; an odor gradient was allowed to form for five minutes before 10 worms were placed in the center of the plate. Chemotaxis was measured after 15 minutes. Chemotaxis index was measured as (worms at odor – worms at control) / total worms.

### Statistics

Given the design of our experiments, we analyzed data in two ways. First, the raw data was analyzed in GRAPHPAD Prism by three-way or two-way ANOVA. When the ANOVA analysis found an effect of or interaction with mutant genotype, we followed up with post-hoc tests using Šidák’s correction that compared each mutant group (fed and food deprived) to the matched WT control animals tested on the same days. (Dunnett’s post hoc tests were used in experiments that had a single WT control group.) This analysis answered the questions, “Do fed mutant animals exit more, less, or about the same as fed WT animals?” and “Do food deprived mutant animals exit more, less, or about the same as food deprived WT animals?” We present this raw data as violin graphs with each raw data point overlaid on the violin graph. For the visual representation of significance in all graphs, we use the following designations: ns, p >= 0.05; *, p < 0.05; **, p < 0.01; ***, p < 0.001; ****, p < 0.0001. The significance results of all two-way and three-way ANOVA interactions are displayed in Table 1; significance results are reported starting as the highest-order interaction and continuing through lower-order interactions and then main effects, if no interaction was significant.

**Table 1:** Analysis of Variance. For each figure, the type of ANOVA is listed, as well as the relevant interactions or main effects. As is standard for the analysis of ANOVAs, the highest order interaction is shown first (note that there are no 3-way interactions for 2-way ANOVAs); if the interaction is not significant, the lower-order interactions are then shown; if no interactions are significant, the main effects are shown. “Mutant genotype” indicates the differences between multiple mutant genotypes when more than one mutant is analyzed in a single figure; “WT control” indicates the difference between each mutant genotype and its respective WT control group when more than one mutant is analyzed in a single figure; “genotype” indicates the differences between WT and a single mutant genotype when only one mutant genotype is analyzed in a figure.

| Figure | Type of ANOVA | 3-way interaction | 2-way interaction | Main Effects |
| --- | --- | --- | --- | --- |
| 2a | 3-way | interaction of mutant genotype x fed condition x WT control, $p < 0.0001$ | n/a | n/a |
| 2b | 2-way | n/a | mutant genotype x WT control, $p < 0.0001$ | n/a |
| 3a | 3-way | mutant genotype x fed condition x WT control, $p = 0.0003$ | n/a | n/a |
| 3b | 2-way | n/a | mutant genotype x WT control, $p < 0.0001$ | n/a |
| 3c | 2-way | n/a | genotype x fed condition, $p < 0.0001$ | n/a |
| 3d | 2-way | n/a | genotype x duration of food deprivation, $p = 0.1094$ | genotype, $p < 0.0001$; duration of food deprivation, $p = 0.0273$ |
| 4a | 3-way | genotype x fed condition x fructose concentration, $p = 0.5291$ | genotype x fed condition, $p = 0.1031$; genotype x fructose concentration, $p = 0.0045$; fed condition x fructose concentration, $p = 0.1518$ | n/a |
| 4b | 2-way | n/a | genotype x fructose concentration, $p = 0.3443$ | fructose concentration, $p = 0.0342$; genotype, $p = 0.0158$ |
| 4c | 2-way | n/a | genotype x fed condition, $p = 0.1774$ | genotype, $p = 0.0011$; fed condition, $p = 0.5771$ |
| 5a | 3-way | mutant genotype x fed condition x WT control, p < 0.0001 | n/a | n/a |
| **5b** | 2-way | n/a | mutant genotype x WT control, p < 0.0001 | n/a |
| **6a** | 3-way | mutant genotype x fed condition x WT control, p = 0.0005 | n/a | n/a |
| **6b** | 2-way | n/a | mutant genotype x WT control, p < 0.0001 | n/a |
| **7a** | 3-way | mutant genotype x fed condition x WT control, p = 0.1428 | mutant genotype x fed condition, p < 0.0001; mutant genotype x WT control, p < 0.0001; fed condition x WT control, p < 0.0001 | n/a |
| **7b** | 2-way | n/a | mutant genotype x WT control, p = 0.0102 | n/a |
| **7c** | 2-way | n/a | genotype x fed condition, p = 0.09162 | genotype, p < 0.0001; fed condition, p < 0.0001 |
| **8a** | 3-way | mutant genotype x fed condition x WT control, p = 0.0509 | mutant genotype x fed condition, p < 0.0001; interaction of mutant genotype x WT control, p < 0.0001; interaction of fed condition x WT control, p < 0.0001 | n/a |
| **8b** | 2-way | n/a | Mutant genotype x WT control, p = 0.0002 | n/a |

However, our research is primarily interested in the question of, “Is the ***effect of food deprivation*** in this mutant larger, smaller, or about the same as in WT animals?” This question is not directly answered by classical two-way or three-way ANOVA with post hoc tests (examples of this are discussed in our results, notably in reference to the behavior of *age-1* and *daf-28* mutant worms). Bootstrapping allows for more complex direct comparisons. We directly tested the effect of food deprivation (i.e. the difference in exiting between fed and food deprived animals in a single genotype) by creating a bootstrapped difference in python (similarly to the bootstrapping of learning integration index in Koemans, Oppitz, *et al*. 2017; Koemans, Kleefstra, *et al*. 2017). Bootstrapping involves repeated random resampling with replacement over many replicates (typically 10,000 replicates). Each replicate makes a new dataset with the same N as the original dataset, resampling to take a random datapoint from the raw dataset to fill each row in the new dataset; for instance, if the raw dataset is “5, 6, 4”, a random resample with replacement could be “4, 6, 6” or “5, 4, 6” (an additional example is shown in **Fig. 1b**). To create a bootstrapped difference, our python script (included as supplemental information) first created a bootstrapped distribution of each raw dataset. After creating 10,000 resampled datasets for each raw dataset, a ‘difference’ dataset was created by calculating the food deprived percent exiting minus the fed percent exiting for 10,000 paired resampled datasets (example in **Fig. 1b**). (Where the sample sizes of the fed and food deprived groups were not equal, we used the smaller N to calculate the difference dataset.) This allowed us to then calculate an average and standard deviation for each of the 10,000 datasets, followed by calculating the mean of the averages and standard deviations. Additionally, we sorted the means of the 10,000 resampled datasets, then removed the 2.5% highest and 2.5% lowest resampled replicate means. We used the remaining maximum and minimum values as the bounds of the 95% confidence interval of the population mean of the effect of food deprivation.

These bootstrapped calculations are robust measures of the effect of food deprivation in each genotype, which then allow us to present targeted statistical and visual analysis. We used the mean average and standard deviation of the bootstrapped difference in GRAPHPAD Prism to test a two-way ANOVA for each data group, followed by post-hoc tests with Šidák’s correction. (For the N of each bootstrapped difference in the statistical analysis, we used the total fed and food deprived N of the raw data minus one, as N – k equals N – 1 when there are two groups, in this case “Fed” and “Food Deprived”). In our figures, we present the bootstrapped mean average of the difference between fed and food deprived for each genotype and WT control group as a mean and 95% confidence interval.

## RESULTS

### Foraging Risk Assay

Our foraging risk assay tests the propensity for worms to exit a ring of an aversive substance, a hyperosmotic fructose solution absorbed into the surface of an agarose plate, in order to approach an attractive food-related odorant (diacetyl). We ran this assay in WT worms after varying lengths of food deprivation (**Fig. 1c**, one-way ANOVA p < 0.001; and **Fig. 1d**, one-way ANOVA p = 0.0003). We found that worms significantly increase their exiting after as little as 30min food deprivation (**Fig. 1c**, p < 0.0001), and continue to exhibit increased exiting compared to fed worms through 3 hours of food deprivation (**Fig. 1c**, p < 0.0001). When comparing the effects of food deprivation of various durations, there was no significant difference between 30 min or 45 min food deprivation when compared to 60 min food deprivation (**Fig. 1d**, respectively p = 0.1768 and p = 0.9882). However, worms that were food deprived for 3 hours exhibited a significantly higher effect of food deprivation than worms that had been food deprived for 1 hour (**Fig. 1d**, p = 0.0166).

### Targeted Screen

We first ran a targeted screen of candidate genes that previous literature suggested might transduce information about the worm’s metabolic state to the neurons involved in making foraging risk decisions (Ao et al. 2004; Apfeld et al. 2004; Murakami et al. 2005; Rottiers and Antebi 2006; Chalasani et al. 2007; Cohen et al. 2009; Dallière et al. 2016). Of the seven gene mutants we tested, three exhibited a strong reduction in the effect of food deprivation (**Fig. 2a** and **2b**; significance for all two-way and three-way ANOVAs is reported in Table 1), while four genes – the AMP-activated protein kinase *aak-2,* the nuclear hormone receptor *daf-12,* the NPY-like peptide *flp-18,* and the glutamate-gated chloride channel *glc-3 –* did not impact foraging risk behavior (**Fig. 2a** and **2b**, p > 0.05). One of the three positive results, the *odr-10* mutant, was an internal positive control that behaved as expected. ODR-10 is an olfactory receptor necessary for sensing the diacetyl odorant used in the multisensory foraging risk assay (Sengupta et al. 1996); thus, we expected the *odr-10* mutant worm to exit similarly to WT animals in the unisensory fructose assay, where no diacetyl was added. This is the case, as *odr-10* mutants in the multisensory foraging risk assay display lower exiting when food deprived (**Fig. 2a**, p < 0.0001) as well as a reduced effect of food deprivation when compared to WT animals (**Fig. 2b**, p < 0.0001). Effectively, the worms do not sense the attractive odorant, and so they do not display the odor-driven enhancement in exiting behavior seen by food deprived worms with normal chemosensory ability.

**Figure 2:**
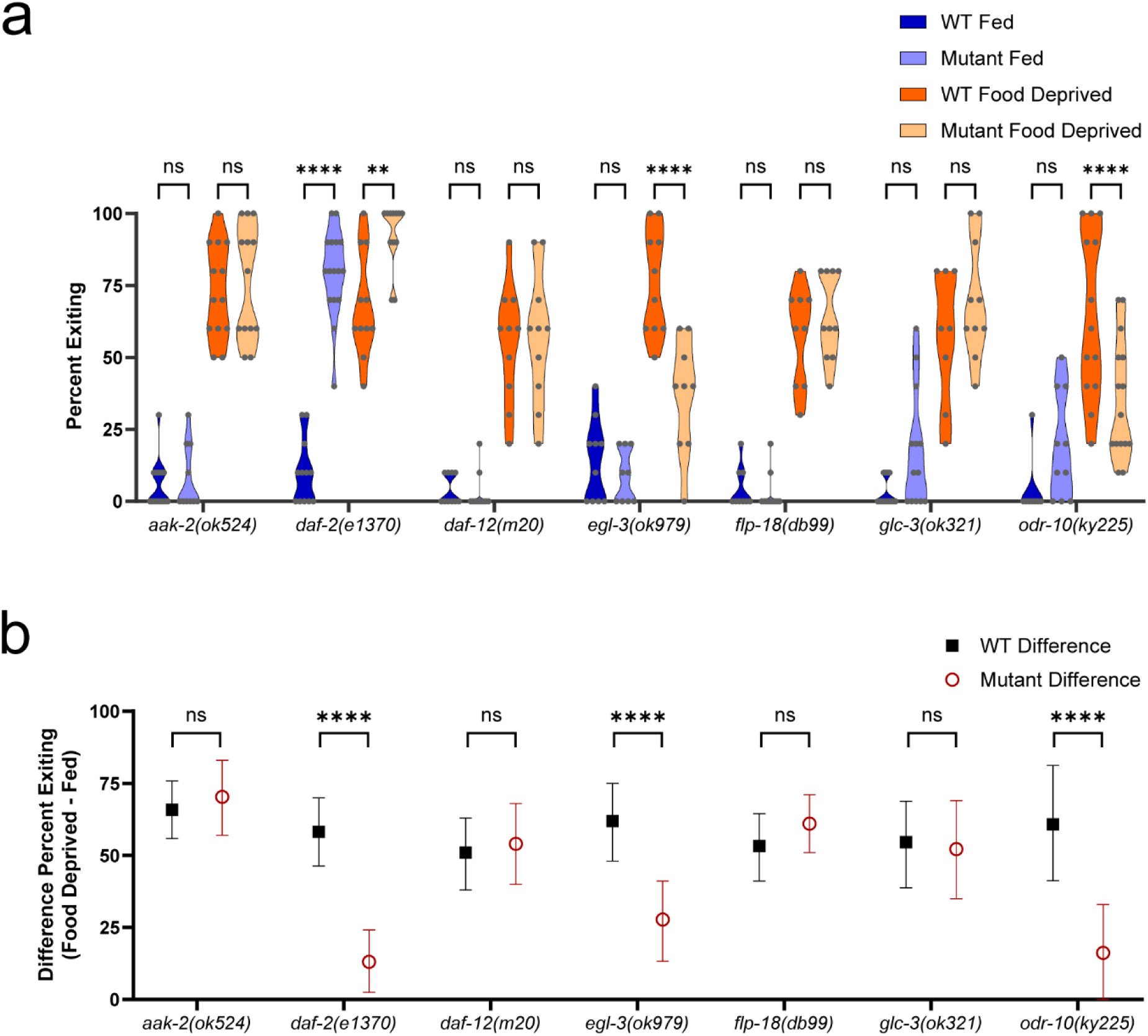
Targeted screen of candidate genes. a) Foraging behavior in the foraging risk assay of fed and food deprived mutant worms and respective WT controls (analyzed by three-way ANOVA, followed by post-hoc tests with Šidák’s correction for WT versus mutant at each condition and genotype). b) Bootstrapped distributions (means and 95% confidence intervals) of the differences between fed and food deprived worms from 2a (analyzed by two-way ANOVA, followed by post-hoc tests with Šidák’s correction for each genotype). Adjusted p-values of post hoc tests: ns, p >= 0.05; **, p < 0.01; ****, p < 0.0001.

The proprotein convertase *egl-3* mutant exhibits foraging behavior that, on a surface level, looks similar to the behavior of the *odr-10* mutant. However, unlike the *odr-10* mutant, the *egl-3* mutant likely has unisensory responses to diacetyl and high-osmolarity fructose that are similar to WT. *egl-3(ok979)* mutant worms have near-normal GCaMP responses to pulses of diacetyl, with perhaps a slight enhancement of habituation and desensitization (Larsch et al. 2015), and the *egl-3(nu349)* mutant has typical responses to hyperosmolarity (Kass et al. 2001). Moreover, it was previously found that *egl-3* mutation reduced a starvation-induced change in backwards movements and tail bending, a microbehavior which may contribute to the changes we see in the foraging risk assay (Thapliyal et al. 2023). Thus, it may be more likely that *egl-3* functions in a hunger signaling pathway, such that loss of the gene results in worms that take lower foraging risks (reduced exiting when food deprived, **Fig. 2a**, p < 0.0001) and exhibit reduced effects of food deprivation (**Fig. 2b**, p < 0.0001).

In contrast to *egl-3*, the *daf-2* IIS receptor gene appears to function as a satiation signal, such that loss of *daf-2* results in worms that exhibit high percent exiting regardless of metabolic state (**Fig. 2a**; higher exiting when well-fed p < 0.0001, higher exiting when food deprived p = 0.0065), with a strong reduction in the effect of food deprivation (**Fig. 2b**, p < 0.0001). DAF-2 is the sole IIS transmembrane receptor in *C. elegans*, and has long been known to play roles in various types of learning where worms avoid an environmental condition associated with food deprivation, such as high salt or altered ambient temperature (Murakami et al. 2005; Kodama et al. 2006; Tomioka et al. 2006; Vellai et al. 2006). Many genes in the IIS pathway in *C. elegans* are orthologous to insulin pathway genes in mammals, suggesting conserved effects of these genes that may translate into mammalian research (Hua et al. 2003). Moreover, it is possible that EGL-3 exerts its influence on foraging behavior via modulation of DAF-2 activity. EGL-3 has been found to process several ILPs which act on the DAF-2 receptor, including INS-3, INS-4, and INS-6 (Hung et al. 2014; Zheng et al. 2018). We thus focused further on the role of the IIS pathway in integrating metabolic state with foraging risk tolerance in the worm.

### Validation of the IIS receptor daf-2

We performed a series of experiments testing for allele-specific effects of *daf-2* and further validating it as a relevant gene in this behavior. First, we tested five additional *daf-2* IIS receptor mutant alleles in the foraging risk assay (**Fig 3a** and **3b**). All of these additional mutant alleles exhibited an increase in exiting when fed (**Fig. 3a**; *daf-2(m41)* p = 0.0001, *daf-2(m596)* p < 0.0001, *daf-2(m579)* p < 0.0001, *daf-2(m577)* p < 0.0001, and *daf-2(e1371)* p < 0.0001), and all but one exhibited a reduced effect of food deprivation (**Fig. 3b**; *daf-2(m41)* p = 0.0338, *daf-2(m596)* p < 0.0001, *daf-2(m579)* p < 0.0001, *daf-2(m577)* p = 0.8501, and *daf-2(e1371)* p < 0.0001). *daf-2* mutant alleles are known to be divisible into two classes: Class I alleles have a more restricted phenotype that only impacts dauer arrest, longevity, and thermotolerance, while Class II alleles have more widespread phenotypic effects, including altered body morphology and brood size, as well as sometimes arresting at embryonic and L1 life stages. It has been hypothesized that Class I alleles may only reduce the ability of *daf-2* to transduce signals to one major downstream effector pathway, while Class II alleles may also impact a second downstream effector pathway (Gems et al. 1998). We found that Class II alleles had overall stronger impacts than Class I alleles in the foraging risk assay. The two alleles which exhibit effects of food deprivation most similar to WT – *m41* exhibiting only a small reduction in the effect of food deprivation (**Fig. 3a**, p = 0.0001), and *m577* exhibiting no reduction in the effect of food deprivation (**Fig. 3b**, p = 0.8501) – are both Class I alleles, while the Class II alleles (*m596, m579,* and *e1370*) all exhibited strong effects. This is consistent with the possibility that multiple downstream *daf-2* effector pathways may be involved.

**Figure 3:**
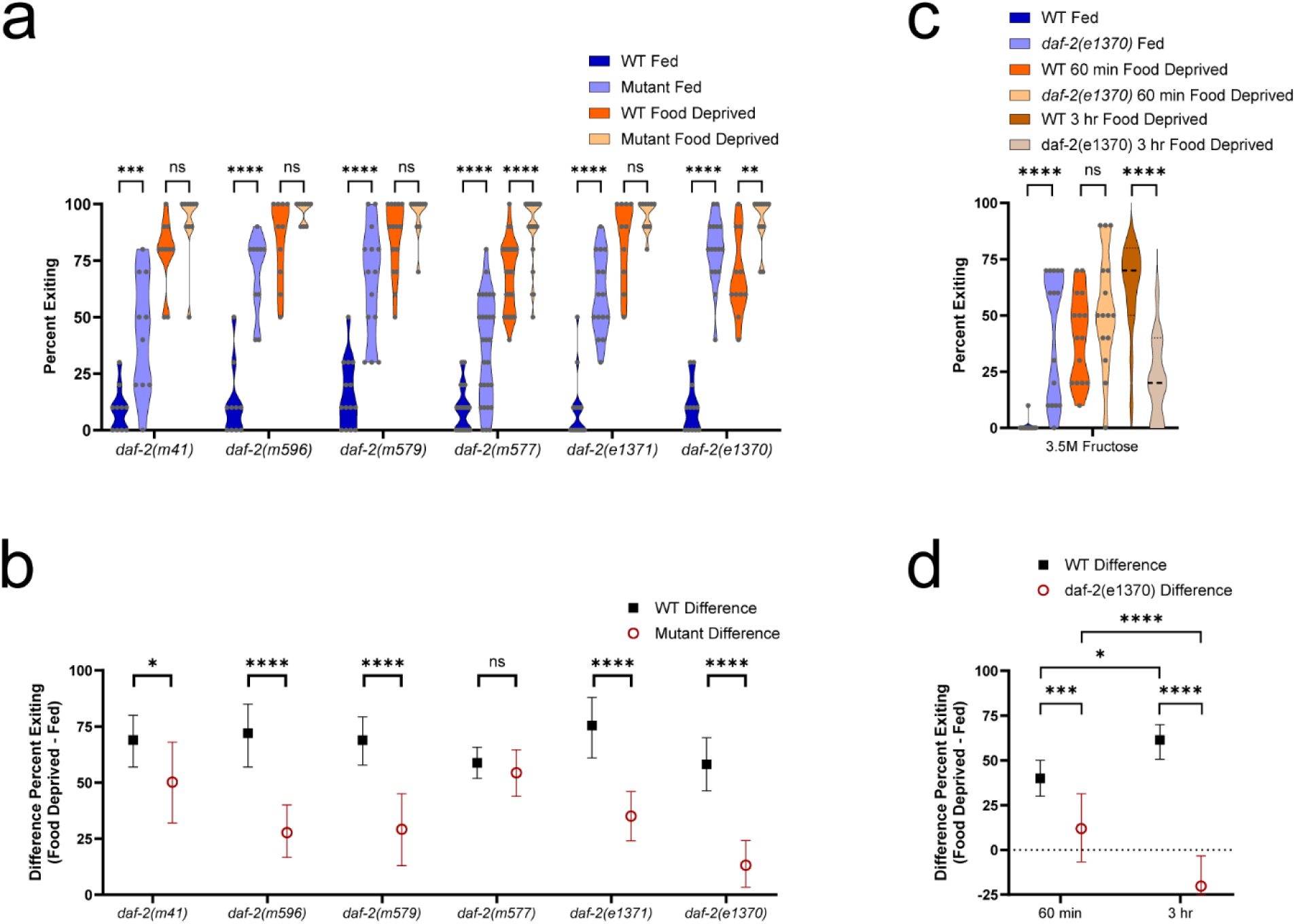
*daf-2* mutant alleles. a) and b) Foraging behavior of worms in the foraging risk assay with different *daf-2* mutant alleles (*daf-2(e1370)* data here is the same as that presented in Fig. 2, shown here for comparison). c) and d) Foraging behavior of *daf-2* mutant worms in the foraging risk assay at increased (3.5M) fructose concentration (using both 60min and 3hr food deprivation). a) and c) show raw data of fed and food deprived mutant worms and respective WT controls (analyzed by three-way ANOVA, followed by post-hoc tests with Šidák’s correction for WT versus mutant at each condition and genotype); b) and d) show the bootstrapped distributions (means and 95% confidence intervals) of the differences between fed and food deprived worms from each respective raw data figure (analyzed by two-way ANOVA, followed by post-hoc tests with Šidák’s correction for each genotype). Adjusted p-values of post hoc tests: ns, p >= 0.05; *, p < 0.05; **, p < 0.01; ****, p < 0.0001.

We next tested the *daf-2* IIS receptor mutant for an effect of food deprivation in a higher-risk version of the foraging risk assay (**Fig. 3c** and **3d**). When a mutant worm shows overall very high exiting behavior, as most *daf-2* mutants do (Fig. **3a** and **3b**), the high baseline behavior may mask an effect of food deprivation. The higher-risk foraging risk assay lowers the percent of worms exiting the circle by increasing the fructose concentration (from a standard 3M to 3.5M) and hence increasing the aversion of worms to exiting. Lowering the overall percent of worms exiting the circle thus allows better observation of any effect of food deprivation. We additionally performed this assay both at our typical (60min) and a longer (3 hour) duration of food deprivation. In the higher-risk foraging assay, *daf-2* mutant worms maintained their increased exiting compared to WT when well fed (**Fig. 3c**, fed WT vs fed *daf-2* mutant, p < 0.0001) but did not display substantially higher exiting than WT when food deprived for 1 hour (**Fig. 3c**, p = 0.2728). At 3 hr food deprivation, *daf-2(e1370)* mutant worms displayed significantly lower exiting than WT worms (**Fig. 3c**, p < 0.0001). Further, we saw that WT worms exhibited greater effect of food deprivation than *daf-2* mutant worms in the 3.5M fructose multisensory foraging risk assay at both 60 min and 3 hr lengths of food deprivation (**Fig. 3d**, p = 0.0005 and p < 0.0001, respectively). While WT worms after 3 hr food deprivation exhibited a stronger effect of food deprivation than after 60 min food deprivation (**Fig. 3d**, p = 0.0127), *daf-2* mutant worms after 3 hr food deprivation exhibited a reduced effect of food deprivation compared to the effect found after 60 min food deprivation (**Fig. 3d**, p < 0.0001). This is unlikely to be due to any sort of reduction in the ability of *daf-2* mutant worms to tolerate starvation, as the *daf-2* mutant is well-known to be exceptionally tolerant of starvation (Baugh and Hu 2020). Together, these results support the conclusion that the higher exiting of fed *daf-2* mutant worms is due to the mutant worms being unable to appropriately distinguish between a fed and 60 min food-deprived metabolic state.

Next, we tested *daf-2* IIS receptor mutant animals in unisensory assays to test whether sensory differences may be driving the smaller effect of food deprivation. As worms in unisensory assays exhibit a much lower effect of food deprivation compared to their behavior in the multisensory foraging risk assay, a lack of ability to sense either hyperosmolarity or diacetyl odor could produce results similar to the results we would expect from a loss of ability to distinguish between fed and food deprived conditions. However, our results indicate that *daf-2* mutant worms are not deficient in their ability to sense hyperosmolarity or the diacetyl odorant (consistent with Matsuura et al. 2007). We tested both WT and *daf-2* mutant animals in a unisensory assay of hyperosmolar avoidance at concentrations of both 3M and 2M fructose (**Fig. 4a** and **4b**). *daf-2* mutants continued to exhibit higher exiting than WT animals when well-fed (**Fig. 4as**, 3M fructose fed WT vs fed *daf-2* mutant, p = 0.0397; 2M fructose fed WT vs fed *daf-2* mutant, p < 0.0001), while also exhibiting the expected higher exiting at 2M fructose than at 3M fructose (**Fig. 4a**). Thus, this assay indicates that the *daf-2* mutant worm ability to sense fructose hyperosmolarity is similar to the ability of a WT worm.

**Figure 4:**
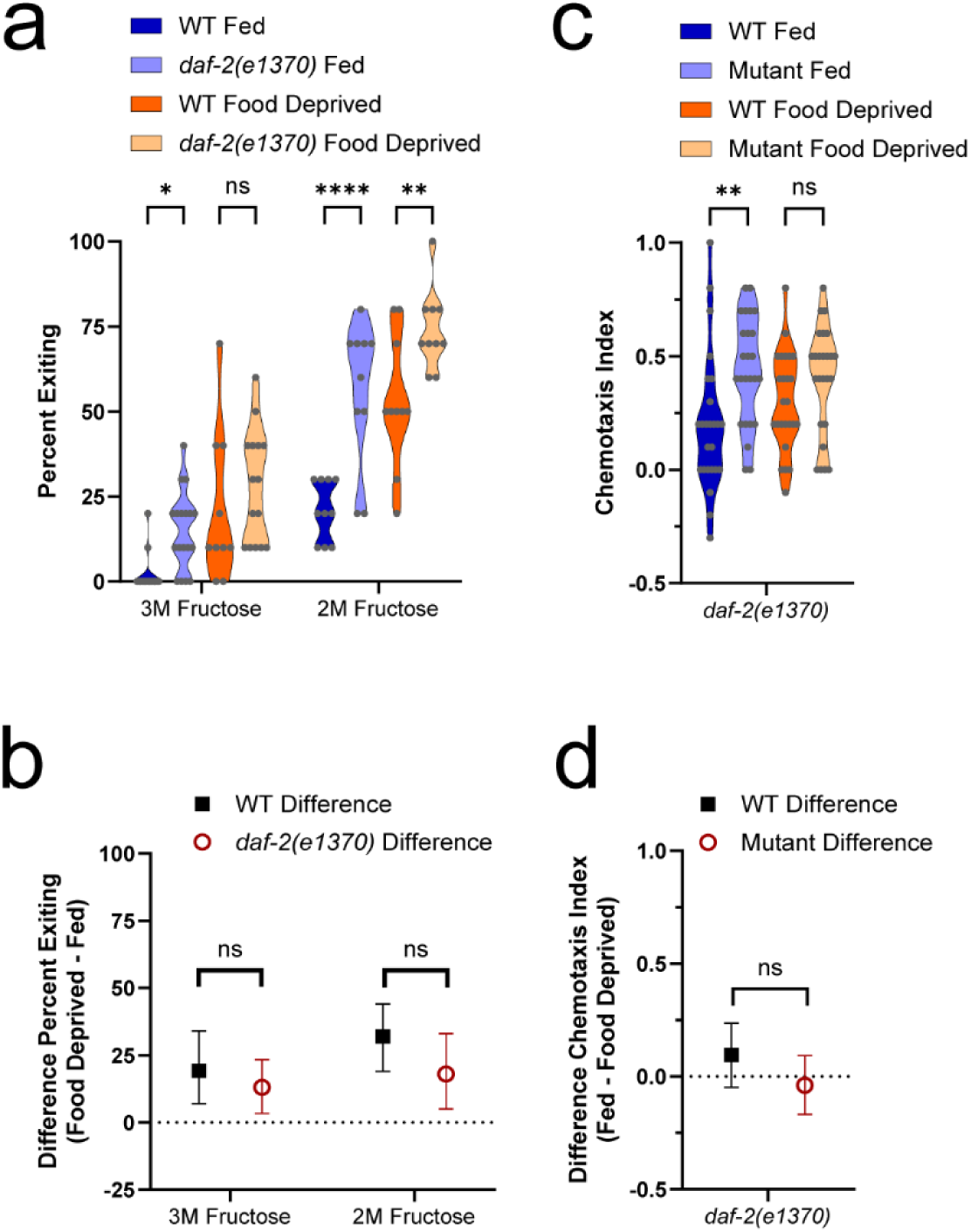
*daf-2* unisensory assays. a) and b) Unisensory hyperosmotic avoidance measured as exiting of the fructose ring by *daf-2* mutants at typical (3M) and reduced (2M) fructose concentration. c) and d) Chemotaxis of *daf-2* mutants to 1µl of 1:10,000 diacetyl. a) and c) show raw data of fed and food deprived mutant worms and respective WT controls (analyzed by three-way ANOVA, followed by post-hoc tests with Šidák’s correction for WT versus mutant at each condition and genotype); b) and d) show the bootstrapped distributions (means and 95% confidence intervals) of the differences between fed and food deprived worms from each respective raw data figure (analyzed by two-way ANOVA, followed by post-hoc tests with Šidák’s correction for each genotype). Adjusted p-values of post hoc tests: ns, p >= 0.05; *, p < 0.05; **, p < 0.01; ****, p < 0.0001.

We also tested both WT and *daf-2* IIS receptor mutant animals in a unisensory assay of chemotaxis to the diacetyl odorant (**Fig. 4c** and **4d**). We found that *daf-2* mutant animals overall exhibited a strong chemotaxis response, while again exhibiting a significant increase in chemotaxis compared to WT animals when well-fed (**Fig. 4c**, WT fed vs. *daf-2* mutant fed, p = 0.0024). This data is both consistent with the hypothesis that the *daf-2* mutant worms are deficient only in their ability to modulate their behavior due to metabolic state (not in their ability to sense diacetyl), and consistent with previous literature finding no deficits in *daf-2(e1370)* chemotaxis (Kauffman et al. 2010).

### Genes of the canonical IIS pathway

We next built on these findings by investigating components of the IIS pathway that are downstream of the *daf-2* IIS receptor (**Fig 5a** and **5b**). Overall, given that our results in *daf-2* mutants indicate that activation of the IIS pathway acts as a satiety signal, we expect that loss-of-function mutants of positive elements of the IIS pathway - *ist-1, age-1, akt-1,* and *akt-2 -* will exhibit higher levels of exiting in the foraging risk assay (i.e. will behave as if they are more hungry), and loss-of-function mutants of negative elements of the IIS pathway - *daf-18* and *daf-16 –* as well as the gain-of-function mutant of the positive element *pdk-1(mg142) -* will exhibit lower levels of exiting (i.e. will behave as if they are less hungry).

**Figure 5:**
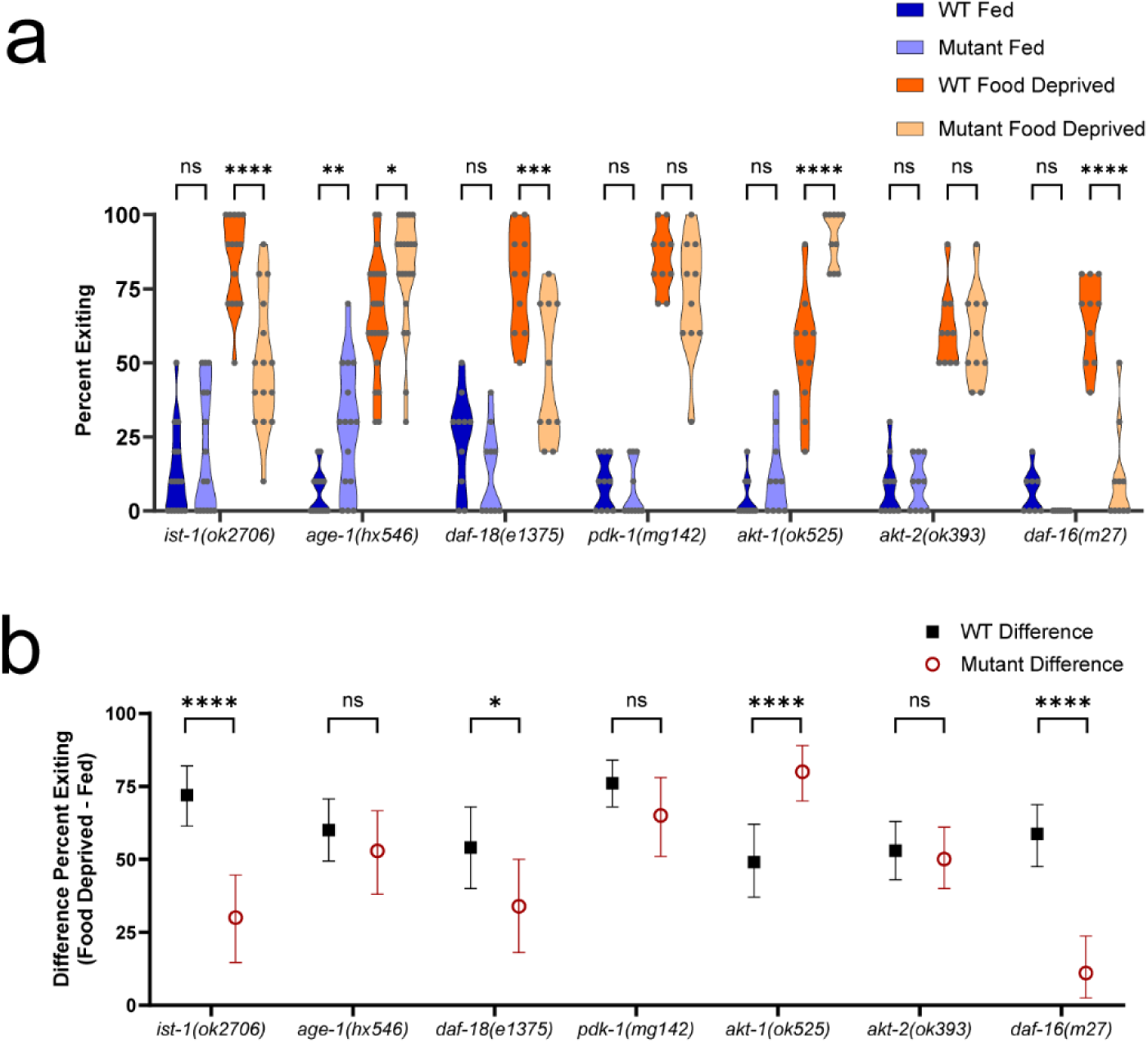
Insulin/IGF-1 signaling pathway mutants. a) Foraging behavior in the foraging risk assay of fed and food deprived mutant worms and respective WT controls (analyzed by three-way ANOVA, followed by post-hoc tests with Šidák’s correction for WT versus mutant at each condition and genotype). b) Bootstrapped distributions (means and 95% confidence intervals) of the differences between fed and food deprived worms from 5a (analyzed by two-way ANOVA, followed by post-hoc tests with Šidák’s correction for each genotype). Adjusted p-values of post hoc tests: ns, p >= 0.05; *, p < 0.05; **, p < 0.01; ***, p < 0.001; ****, p < 0.0001.

Contrary to our expectations, the *ist-1* mutant exhibited overall lower exiting when food deprived (**Fig. 5a**, p < 0.0001) with a greatly reduced effect of food deprivation (**Fig. 5b**, p < 0.0001). This lower exiting when food deprived could be due to IST-1 acting via non-canonical IIS pathways, rather than simply as a positive element in the canonical IIS pathway. As we predicted given the canonical IIS pathway, *age-1* mutants exhibited higher exiting than WT worms, both when fed and when food deprived (**Fig. 5as**, fed WT vs fed *age-1* mutant p = 0.0031, food deprived WT vs food deprived *age-1* mutant p = 0.0330). However, *age-1* mutants exhibited a similar effect of food deprivation compared to WT worms (**Fig. 5b**, p = 0.6621). As *age-1(hx546)* mutants have reduced AGE-1 activity, not a complete loss of function (Morris et al. 1996), it is possible that the *age-1(hx546)* mutants retained sufficient AGE-1 function to still respond to changes in metabolic state. Alternatively, these results could suggest that while the effect of AGE-1 activity is to lower high-risk foraging activities overall, AGE-1 activity may not be required for metabolic state to alter the worm’s behavior.

The behavior of *age-1* mutant worms additionally illustrates one scenario where including bootstrapped effect data is superior to only using traditional statistical tests. Our bootstrapped analysis allowed us to determine that, despite differences between the raw exiting in both fed and food deprived groups, the effect of food deprivation on exiting in *age-1* mutants did not differ from that of WT worms (**Fig. 5b**, p = 0.6621). Traditional ANOVA statistical tests with direct posthoc comparisons of raw data will not differentiate between data such as our *age-1* mutant worm data, and data where a difference in the same direction in both food and fed deprived worms does impact the effect of food deprivation (such as the behavior of *daf-2* receptor mutant worms in **Fig. 2a** and **2b**).

The remaining kinases and phosphatases that we tested in the IIS pathway also exhibited results that were overall consistent with our hypothesis of IIS pathway activation acting as a satiety signal. The *daf-18* (PTEN) mutant exhibited reduced exiting when food deprived (**Fig. 5a**, p = 0.0003), as well as a reduction in the effect of food deprivation (**Fig. 5b**, p = 0.0179). While *akt-1* mutant worms exhibited the expected increased exiting when food deprived (**Fig. 5a**, p < 0.0001), they were the only mutant worm we tested that exhibited a significantly *increased* effect of food deprivation compared to WT worms (**Fig. 5b**, p < 0.0001). This may suggest the involvement of a non-canonical pathway, as the worm is able to allow metabolic state to modulate its foraging behavior without the influence of *akt-1.* By contrast, the *akt-2* mutant worm behaved similarly to WT (**Fig. 5a** and **5bs**, p > 0.05). These results are likely at least in part because AKT-1 and AKT-2 have some degree of redundancy in worms (Paradis and Ruvkun 1998); additionally, it is possible that *akt-1* plays a larger role than *akt-2* in neurons involved in modulating foraging behavior, as a significant subset of neurons express *akt-1* but not *akt-2* (Taylor et al. 2021). Finally, *daf-16* FOXO transcription factor mutant worms exhibited reduced exiting when food deprived compared to WT worms (**Fig. 5a**, p < 0.0001) and a reduced effect of food deprivation (**Fig. 5b**, p < 0.0001). Together these results support the canonical IIS pathway, from the receptor *daf-2* to the transcription factor *daf-16*, as a necessary modulator of foraging risk in *C. elegans*.

We additionally tested several ILP (insulin-like peptide) mutants (**Fig 6a** and **6b**). *C. elegans* has 40 ILPs, which can have wide variation in their expression (Ritter et al. 2013). Some, like INS-1 and INS-7, act as antagonists, rather than agonists, of the DAF-2 receptor (Pierce et al. 2001; Chen et al. 2013). As our hypothesis is that activation of the IIS pathway acts as a satiety signal (to reduce high-risk foraging behavior, and hence reduce exiting in the foraging risk assay), we expected that mutants of ILPs that act as agonists would exhibit overall increased exiting, and mutants of ILPs that act as antagonists would exhibit overall reduced exiting.

**Figure 6:**
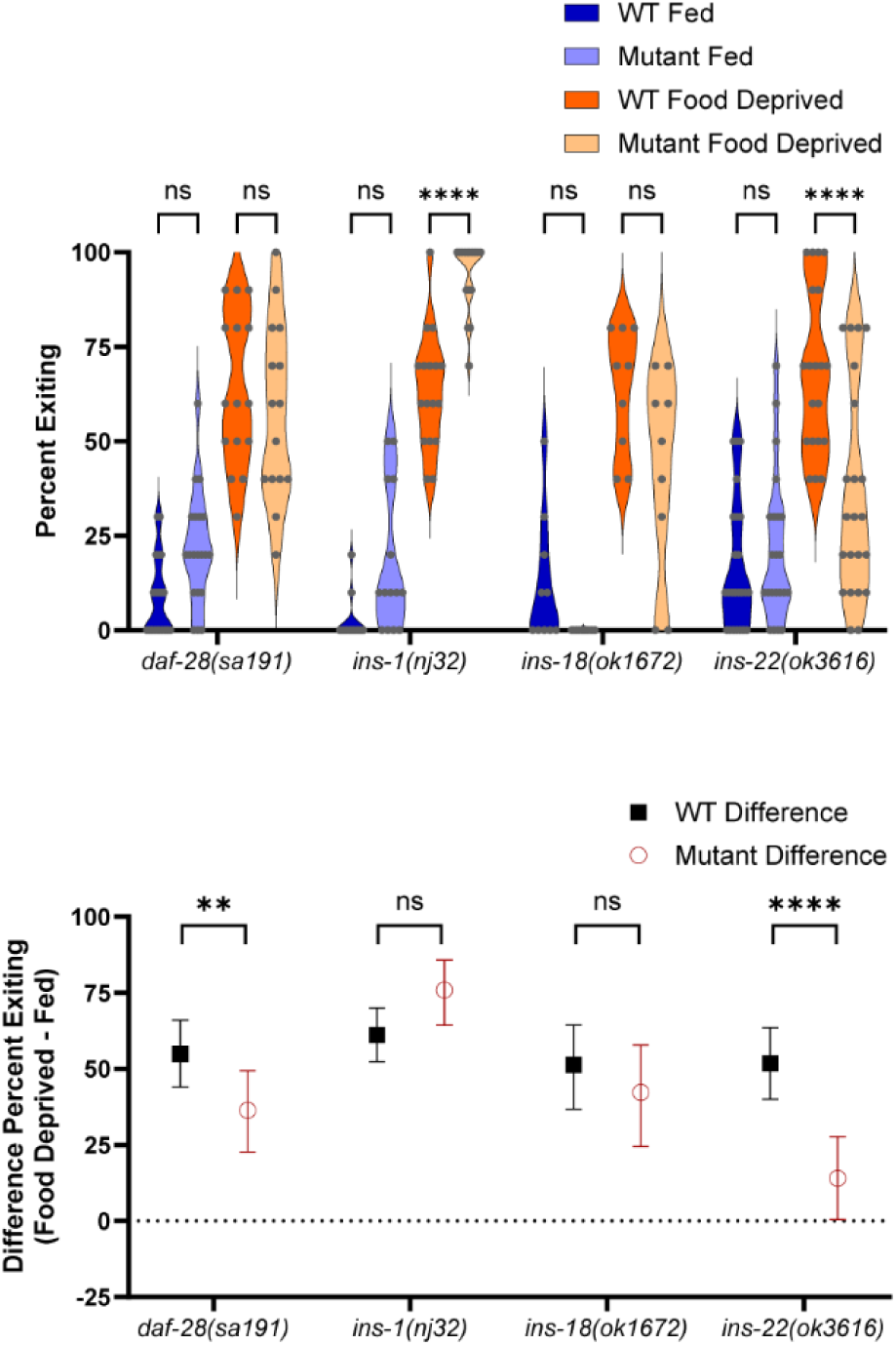
Insulin-like peptide mutants. a) Foraging behavior in the foraging risk assay of fed and food deprived mutant worms and respective WT controls (analyzed by three-way ANOVA, followed by post-hoc tests with Šidák’s correction for WT versus mutant at each condition and genotype). b) Bootstrapped distributions (means and 95% confidence intervals) of the differences between fed and food deprived worms from 6a (analyzed by two-way ANOVA, followed by post-hoc tests with Šidák’s correction for each genotype). Adjusted p-values of post hoc tests: ns, p >= 0.05; *, p < 0.05; **, p < 0.01; ***, p < 0.001; ****, p < 0.0001.

We found that a *daf-28* ILP mutant, which is characterized as an agonist of the IIS pathway (Zheng et al. 2018), did not differ significantly from WT animals in either fed or food deprived conditions (**Fig. 6as**, p > 0.05). However, the *daf-28* mutant did exhibit a reduction in the effect of food deprivation (**Fig. 6b**, p = 0.0076). Notably, the behavior of *daf-28* mutants is an excellent example wherein small differences in the opposite direction in both fed and food-deprived groups result in no direct difference between the paired groups following classical ANOVA and posthoc analysis, yet the bootstrapped analysis of the effect of food deprivation reveals an overall change. Slightly lower food deprived exiting and slightly higher fed exiting are not significant differences alone, but together they add up to a significantly smaller effect of food deprivation when bootstrapped effect analysis is conducted.

By contrast, *ins-1* mutant worms exhibited an increase in exiting when food deprived (**Fig. 6a**, p < 0.0001), but did not differ significantly from WT in the effect of food deprivation (**Fig. 6b**, p = 0.0848). This mild effect is consistent with previous work on *ins-1*, which has shown that *ins-1* interaction with the IIS pathway is complex; it may act as an agonist or antagonist of the IIS pathway, depending on the context (Pierce et al. 2001; Kodama et al. 2006; Tomioka et al. 2006; Zheng et al. 2018). *ins-18* mutant worms behaved similarly to WT worms both when fed and food deprived (**Fig. 6as**, p > 0.05) and in the effect of food deprivation (**Fig. 6b**, p = 0.7083). Finally, the *ins-22* mutant worm (which Zheng *et al*. 2018 found to be an antagonist) exhibited both a reduction in exiting when food deprived (**Fig. 6a**, p < 0.0001), as well as a strong reduction in the effect of food deprivation (**Fig. 6b**, p < 0.0001). These results further support the IIS pathway as a modulator of risky foraging behavior and suggest complexity in the ILPs that may transmit signals about metabolic state. The ILP *ins-22* is clearly necessary for the appropriate increase in exiting when food deprived, but other ILPs, such as *daf-28* and *ins-1*, may also modulate risky foraging behavior in more subtle ways.

### Validation of the FOXO transcription factor daf-16

For further validation of *daf-16* (the FOXO transcription factor primarily regulated by the IIS pathway) in transducing metabolic state to foraging behavior, we tested three more *daf-16* mutant alleles (**Fig 7a** and **7b**). All of these *daf-16* mutants exhibited a decrease in exiting when fed (**Fig. 7as**; *daf-16(mu86)* p < 0.0001, *daf-16(m26)* p = 0.0003, *daf-16(mgDf50)* p < 0.0001), and all exhibited a reduced effect of food deprivation (**Fig. 7bs**; *daf-16(mu86)* p < 0.0001, *daf-16(m26)* p = 0.0001, *daf-16(mgDf50)* p < 0.0001).

**Figure 7:**
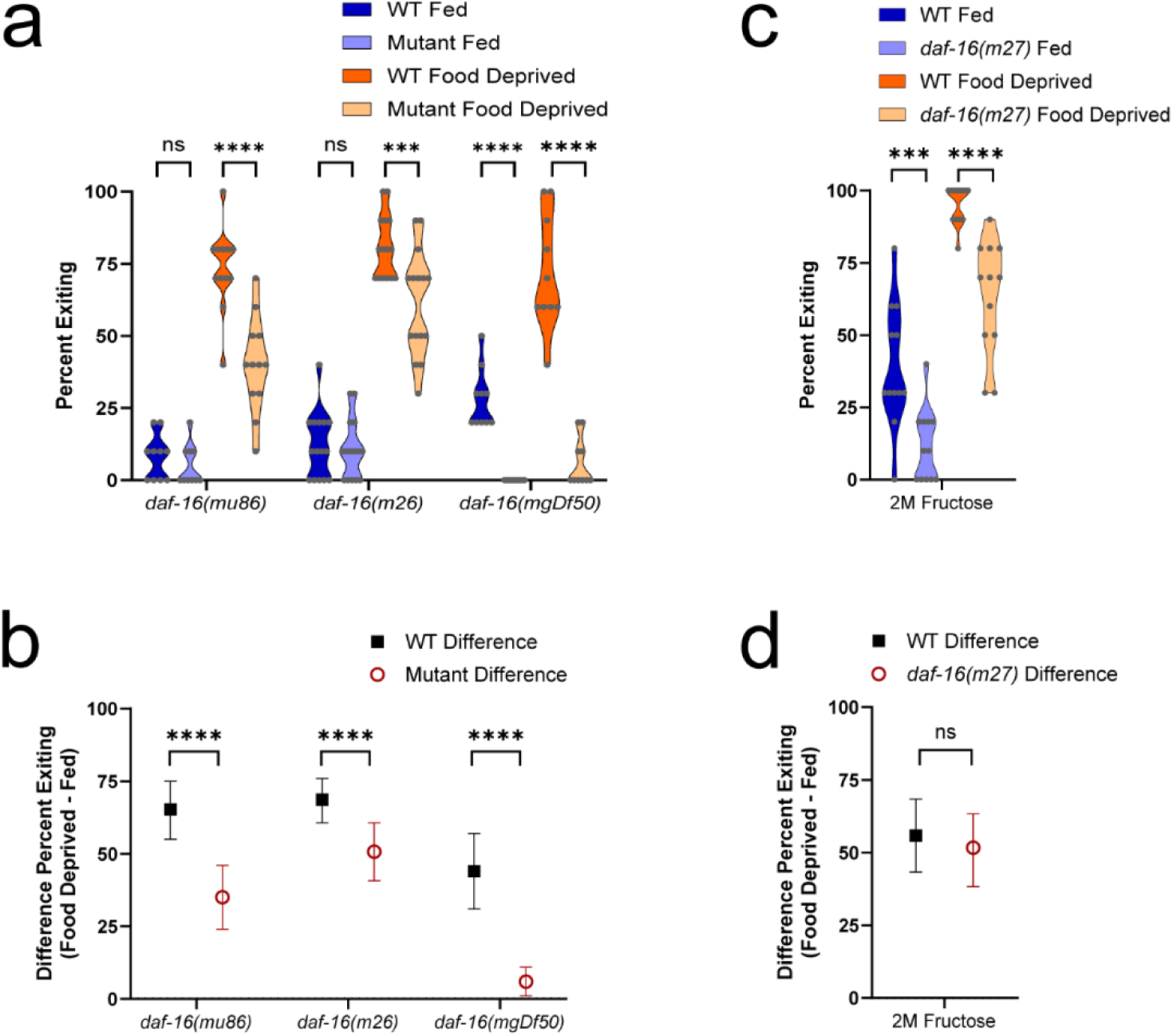
*daf-16* mutant alleles. a) and b) Foraging behavior in the foraging risk assay of worms with different *daf-16* mutant alleles. c) and d) Foraging behavior of *daf-16* mutant worms in the foraging risk assay at reduced (2M) fructose concentration. a) and c) show raw data of fed and food deprived mutant worms and respective WT controls (analyzed by three-way (a) or two-way (c) ANOVA, followed by post-hoc tests with Šidák’s correction for WT versus mutant at each condition and genotype); b) and d) show the bootstrapped distributions (means and 95% confidence intervals) of the differences between fed and food deprived worms from each respective raw data figure ((b) analyzed by two-way ANOVA followed by post-hoc tests with Šidák’s correction for each genotype, (d) analyzed by Student’s t-test). Adjusted p-values of post hoc tests: ns, p >= 0.05; ***, p < 0.001; ****, p < 0.0001.

We next tested the *daf-16* FOXO transcription factor mutant for an effect of food deprivation in a lower-risk version of the foraging risk assay (Fig. **7c** and **7d**). When a mutant worm shows overall very low exiting behavior, as most *daf-16* mutants do (Fig. **7a** and **7b**), the low baseline behavior may mask an effect of food deprivation. The lower-risk foraging risk assay increases the percent of worms exiting the circle by decreasing the fructose concentration (from a standard 3M to 2M) and hence decreasing the aversion of worms to exiting. Raising the overall percent of worms exiting the circle thus allows better observation of any effect of food deprivation. At the lower 2M fructose concentration, *daf-16* mutant worms maintained their overall lower exiting when food deprived (**Fig. 7c**, p < 0.0001) but now also exhibited lower exiting when well fed (**Fig. 7c**, p = 0.0003), resulting in a similar effect of food deprivation compared to WT animals (**Fig. 7d**, Student’s t-test, p = 0.4956). This suggests that, while DAF-16 transcription factor activity increases exiting and thus high-risk foraging behaviors, *daf-16* (unlike the IIS receptor *daf-2*) is not necessary for the worm to distinguish between a fed and food deprived metabolic state. These results further highlight the possibility that the DAF-2 receptor-mediated effect of food deprivation on exiting may be transduced via a non-canonical pathway (i.e., not solely via modulation of DAF-16 transcription factor activity).

### Tests of a non-canonical pathway

To further test the hypothesis that the IIS receptor DAF-2 acts via a non-canonical pathway to modulate foraging risk in response to food deprivation, we tested double mutants of *daf-2* and *daf-16* (**Fig. 8a** and **8b**). If DAF-2 acts solely via the canonical pathway that culminates by inhibiting DAF-16 transcriptional activity, then we would expect a *daf-2; daf-16* double mutant to have a behavioral phenotype very similar to the *daf-16* single mutant (e.g., reduced exiting, especially when food deprived). However, we found that two independent *daf-2; daf-16* double mutants both exhibited a phenotype that was more similar to the *daf-2* IIS receptor single mutant than the *daf-16* FOXO transcription factor single mutant. *daf-2; daf-16* double mutant worms exhibited increased exiting when fed, but lacked a large or significant decrease in exiting when food deprived (**Fig. 8as**, da*f-16(m26); daf-2(e1370)* fed vs WT fed p < 0.0001; *daf-16(mgDf47); daf-2(e1370)* fed vs WT fed p = 0.0021; *daf-16(m26); daf-2(e1370)* food deprived vs WT food deprived p = 0.9892; *daf-16(mgDf47); daf-2(e1370)* food deprived vs WT food deprived p = 0.0668). The *daf-2; daf-16* double mutant worms additionally exhibited a strong reduction in the effect of food deprivation (**Fig. 8bs**, *daf-16(m26); daf-2(e1370)* vs WT p < 0.0001; *daf-16(mgDf47); daf-2(e1370)* vs WT p < 0.0001).

**Figure 8:**
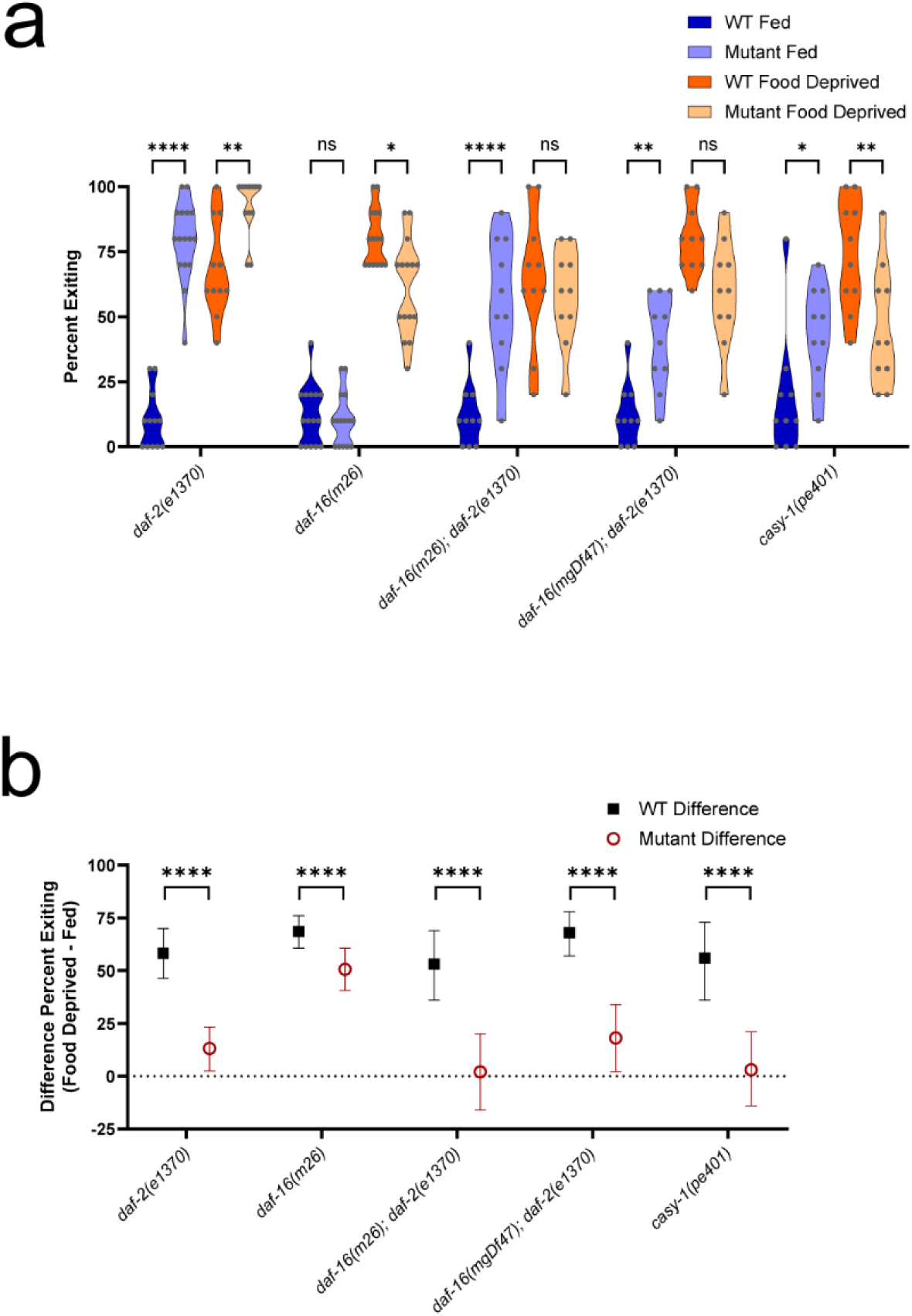
*daf-16; daf-2* double mutants and *casy-1* mutant. a) Raw data of fed and food deprived mutant worms and respective WT controls in the foraging risk assay (analyzed by two-way ANOVA, followed by post-hoc tests with Šidák’s correction for WT versus mutant at each condition and genotype). (Single mutants *daf-2(e1370)* and *daf-16(m26)* are the same data as shown in Fig. 2a and Fig. 5a, respectively, shown here for comparison.) b) Bootstrapped distributions (means and 95% confidence intervals) of the differences between fed and food deprived worms from the raw data in 6a (analyzed by two-way ANOVA, followed by post-hoc tests with Šidák’s correction). Adjusted p-values of post hoc tests: ns, p >= 0.05; *, p < 0.05; **, p < 0.01; ****, p < 0.0001.

We then tested *casy-1* as a potential mediator of a non-canonical *daf-2* pathway in the foraging risk assay. The *daf-2* IIS receptor gene in worms is expressed primarily as two isoforms, *daf-2a* and *daf-2c*. These isoforms differ only in that *daf-2c* includes the ligand binding domain exon 11.5 (Ohno et al. 2014). Previous research has found that inclusion of exon 11.5 allows DAF-2c to bind to CASY-1, which then binds to a kinesin-1 complex. DAF-2c is trafficked to the axon following food deprivation, where it may mediate synaptic activity; its presence is necessary for salt avoidance learning following food deprivation (Ohno et al. 2014). *casy-1* mutants have been previously found to show defects in learning (Hoerndli et al. 2009) and sensory integration (Ikeda et al. 2008), while displaying typical naïve chemotaxis and hyperosmotic avoidance (Ikeda et al. 2008; Hoerndli et al. 2009). In our foraging risk assay, *casy-1* mutant worms exhibited both increased exiting when fed (**Fig. 8a**, p = 0.0134) and decreased exiting when food deprived (**Fig. 8a**, p = 0.0034), leading to the strongest reduction in the effect of food deprivation that we observed in any single mutant, with a near-zero effect of food deprivation on exiting (**Fig. 8b**, *casy-1* effect of food deprivation vs WT effect of food deprivation, p < 0.0001). This further supports the role of a non-canonical *daf-2* pathway in mediating the effects of food deprivation on high-risk foraging behaviors and suggests a new avenue of research regarding the exact mechanisms by which a CASY-1 mediated DAF-2 signal may modulate behavior.

## DISCUSSION

Together, these results suggest a complex, two-pronged pathway by which the IIS receptor may mediate foraging risk behaviors in response to food deprivation. When metabolic state is satiated (worms are well-fed), overall expression of agonistic ILPs is high (and expression of antagonistic ILPs is low) (Chen and Baugh 2014). High levels of agonistic ILPs leads to activation of the IIS receptor DAF-2, which inhibits the FOXO transcription factor DAF-16 and thereby reduces high-risk foraging behaviors. Additionally, when worms are well-fed, the DAF-2c isoform of the IIS receptor is primarily localized to the cell body (Ohno et al. 2014), which reduces DAF-2c activity in axons and may also reduce high-risk foraging behaviors. When food deprivation alters metabolic state, IIS activation is reduced. Food deprivation then increases CASY-1 trafficking of DAF-2c to axons (Ohno et al. 2014), which may increase high-risk foraging behaviors. Additionally, the disinhibition of DAF-16 transcriptional activity leads to an increase in exiting, further strengthening the worm’s response to food deprivation.

We consider it an intriguing possibility that the canonical and non-canonical DAF-2 pathways may in part function to modulate the worm’s response to food deprivation via different time scales. Multiple papers have reported effects of starvation that differ depending on the length of starvation (Murakami et al. 2005; Ghosh et al. 2016), and previous observations from our lab and others have demonstrated changes in foraging strategy in as little in 30 minutes (Hills et al. 2004; Ghosh et al. 2016). However, changes in cellular localization of the FOXO transcription factor DAF-16 due to food deprivation are generally thought to be slow, with most studies observing an effect after several hours to a day or more (Henderson and Johnson 2001; Weinkove et al. 2006), and no studies that we could find demonstrating a change in cellular localization in less than one hour. Notably, one study of daf-16::GFP cellular localization observed nuclear localization after 75 minutes of food deprivation, while they did not see clear nuclear localization after 60 minutes of food deprivation (Henderson and Johnson 2001).

Moreover, transcription factor mediated effects take even more time following the nuclear translocation of the transcription factor, as genes that may function as downstream mediators must be transcribed, and usually also translated, before altering the worm’s behavior. In contrast to the longer timescales of transcription factor activity, the speed of fast axonal transport and length of *C. elegans* axons suggest proteins can be transported from the soma to synapses in 15-30 minutes (Wu et al. 2007; Niwa 2017; Feng et al. 2018; Lipton et al. 2018; Wu et al. 2024), and activation of kinase pathways present at the synapse likely occurs in seconds to minutes (Murphy et al. 1994; Blazek et al. 2015). Thus, it is possible that the DAF-2c / CASY-1 pathway is an early modulator of foraging risk following short-term food deprivation (i.e., in the 15-60 minute time range), while the DAF-2 / DAF-16 pathway is a mechanism that may be slower to act but may also be more strongly maintained over time. In support of this time-separated hypothesis, the food-deprivation-driven salt learning seen in Tomioka et al. 2006 (which used 1 hour of food deprivation) required the *daf-2* IIS receptor but did not seem to be affected by loss of the FOXO transcription factor *daf-16.* A two-pronged foraging modulation strategy over multiple time scales may be especially relevant for an organism like *C. elegans* that has a boom-and-bust population growth strategy. For these worms, very low-risk foraging is ideal during the “boom” phase when food is abundant, but a strongly risk-tolerant foraging strategy that is both timely and sustained is imperative for animals to disperse and some to eventually find more food during the “bust” phase, when a food source has been depleted (Frézal and Félix 2015).

These differences in time scale may in part explain some of the differences between our results, and the results obtained in a recent paper that also used a multisensory foraging risk assay (Matty et al. 2022). Matty et al. 2022 made important contributions to our understanding of foraging regulation in the worm, notably identifying HLH-30 and MML-1 as transcription factors that modulate foraging risk behavior and likely modulate expression of ILPs. Similarly to our assay, they separated worms from a food-related (diacetyl) odor using an aversive barrier (copper sulfate) and counted animals that crossed the barrier vs. animals that did not cross the barrier. Notably, both copper and high-osmolarity fructose are primarily sensed by the ASH neuron (Hilliard et al. 2005); the neural circuits for both assays are likely to be similar. However, while we used an assay test time of 15 minutes and saw changes with as little as thirty minutes of food deprivation, their assay used a test time of 45 minutes and a typical food deprivation duration of 3 hours (Matty et al. 2022). As worms are not fed during the assay test, and the time at which we see alterations in behavior is (including the assay test time) 45 minutes past the initiation of food deprivation, this means that ‘well-fed’ worms in a 45 minute test assay are likely to behave as if they are slightly food deprived (solely by any short-term mechanisms) by the end of the assay. Using the 45 minute test, they did not find a change in foraging behavior at 1 hour food deprivation, but did see a difference at 2 hours (Matty et al. 2022). Thus, we argue that our foraging risk assay using a 15-minute test period and 30- to 60-minutes of food deprivation is primarily testing short-term mechanisms of food-deprivation-modulated risky foraging behavior. By contrast, their foraging risk assay using a 45-minute test period may be testing long-term mechanisms of food-deprivation-modulated risky foraging behavior, which may not begin until 2-3 hours after the initiation of food deprivation.

These results suggest areas of further research into how the insulin signaling pathway in the brain may modulate mammalian food-seeking behaviors and dietary choices. Strong evidence suggests neuronal insulin is an important signaling pathway. Insulin in the mammalian CNS is known to play a role in feeding behavior (Baldini and Phelan 2019), metabolic diseases (Porte et al. 2005), Alzheimer’s disease and other types of neurodegeneration (Kellar and Craft 2020; Alberini 2023), and learning and memory (Spinelli et al. 2019). The isoform of the IIS receptor trafficked by CASY-1 (DAF-2c) has similarities to the mammalian insulin receptor B isoform (Ohno et al. 2014). While the insulin receptor B isoform is not found in mammalian neurons (Kenner et al. 1995), previous research has shown that the insulin receptor can localize to rat synapses (Abbott et al. 1999). It is not yet fully clear in which cells and under which circumstances the mammalian insulin receptor may be trafficked to synapses, which cellular pathways may be modulated at those synapses, and how synaptic effects of the insulin receptor may differ in timescale and physiological effect from actions of the insulin receptor in the cell soma or nucleus. Our research, however, suggests the possibility that an improved understanding of these questions may provide insights into the neural control of mammalian foraging behavior.

## Supporting information

BootstrappingJupyterNotebook

## ACKNOWLEDGEMENTS

We thank Jess Wu, Sanjana Bandi, and Vivian Wang for conducting troubleshooting experiments. This work was supported by an R35 grant from the National Institute of General Medical Sciences (NIGMS) (to M.N.N.).

