## Supplementary material for "Metabolic state modulates risky foraging behavior via multiple branches of the insulin/IGF-1-like pathway in *C. elegans*": BootstrappingJupyterNotebook: PriceEtAlBootstrapping.html


In [ ]:

```
#Write in your data file, and check that it is the correct file. Remember the formatting:
# 1) It must be a .csv file
# 2) Each group is a single column
# 3) The first row is a descriptive name for each group, with no spaces or special characters
# 4) The order of the groups is such that the first column is a "naive" or "untreated",
#    followed by its respective "trained" or "treated" group; then repeat this for all groups, so that
#    all odd columns are "naive" or "untreated" and all even columns are "trained" or "treated"

import numpy as np
RawData = np.genfromtxt('DevVar.csv', delimiter=',', dtype=float, names=True)
print("This will list the groups")
print(RawData.dtype)
print("This will show the first row of the data for all groups")
print(RawData[0])
```

In [ ]:

```
#This will tell the notebook which groups should be paired with which other groups, and create a name for each pair
#that we will use to name the paired data. Here, each row is a different pair; the first column is the "untreated"
#group name, the second column is the "treated" group name, and the third column is the "vs" pair name.

RawNames = list(RawData.dtype.fields)
NumberPairs = int((len(RawNames))/2)
NumberRawGroups = int((len(RawNames)))
ArrNames = np.asarray(RawNames)
DataNames = ArrNames.reshape(NumberPairs,2)

PairNames = np.empty([NumberPairs,], dtype=object)
for i in range(NumberPairs):
    PairNames[i,] = (DataNames[i, 0] + "_VS_" + DataNames[i,1])
    
AllDataNames = np.column_stack((DataNames, PairNames))

print("This will list all of the data groups paired together, plus a third name for the paired group")
print(AllDataNames)
```

In [ ]:

```
#Here we will specify which group we generally expect to be higher.
#This will let us create difference statistics that are normally positive (but could be negative!)
#If the "untreated" wildtype control is usually a  higher number, put 0
#If the "treated" wildtype control is usually a higher number, put 1

BaseDiff = 1
```

In [ ]:

```
#Here we will get the number of data points in each group. The array for length is set up much like the array for
#group names; so the first column is the "Control" group, the second is its paired "Treatment" group, and the third
#is the smallest N of the two (so arrays can be directly compared). Here the fourth column is the N that
#we will use for degrees of freedom (so the sum of the paired groups minus 1).

LenArr = np.empty([NumberPairs,4])

for i in range(NumberPairs):
    LenArr[i,0] = int(np.count_nonzero(~np.isnan(RawData[:][AllDataNames[i,0]])))
    LenArr[i,1] = int(np.count_nonzero(~np.isnan(RawData[:][AllDataNames[i,1]])))
    LenArr[i,2] = int(np.minimum(LenArr[i,0], LenArr[i,1]))
    LenArr[i,3] = (LenArr[i,0] + LenArr[i,1]) - 1
    
print(LenArr)
```

In [ ]:

```
#Next, we'll store array names for the bootstrapped samples for each group, and for an array we will use for the 
#summary data for each pair.

BootNames = np.empty([NumberPairs,5], dtype=object)

for i in range(NumberPairs):
    BootNames[i,0] = "Boot" + AllDataNames[i,0]
    BootNames[i,1] = "Boot" + AllDataNames[i,1]
    BootNames[i,2] = "Boot" + AllDataNames[i,2]
    BootNames[i,3] = "SumBoot" + AllDataNames[i,2]
    BootNames[i,4] = "Stats" + AllDataNames[i,2]

print(BootNames)
```

In [ ]:

```
#Next we will create a bootstrapped sample of 10,000 re-sampled datasets for each data group.
#Our sample size will be the smaller N of each pair, since we will shortly be computing these pairs together.
#The output will print the raw data for each group, and then the group name and "done" as it finishes each one.

Sample = 10000

for i in range(NumberPairs):
    TempGroup = AllDataNames[i,0]
    TempN = int(LenArr[i,0])
    print("The raw data for " + TempGroup + " is ")
    print(RawData[0:TempN][TempGroup])
    BootNames[i,0] = np.random.choice(RawData[0:TempN][TempGroup], size=[Sample, int(LenArr[i,2])])
    print(BootNames[i,0])
    print("done! "+TempGroup)

for i in range(NumberPairs):
    TempGroup = AllDataNames[i,1]
    TempN = int(LenArr[i,1])
    print("The raw data for " + TempGroup + " is ")
    print(RawData[0:TempN][TempGroup])
    BootNames[i,1] = np.random.choice(RawData[0:TempN][TempGroup], size=[Sample, int(LenArr[i,2])])
    print(BootNames[i,1])
    print("done! "+TempGroup)
```

In [ ]:

```
#Here, we will create each paired bootstrap group! This will take each set of bootstrapped data and find the
#difference with its paired bootstrapped group.

if BaseDiff == 0:
    print("Bootstrapping pairs by calculating control minus treated")
else:
    print("Bootstrapping pairs by calculating treated minus control")

for i in range(NumberPairs):
    if BaseDiff == 0:
        BootNames[i,2] = BootNames[i,0] - BootNames[i,1]
        print(BootNames[i,2])
        print(BootNames[i,2].shape)
        print("done!")
    else:
        BootNames[i,2] = BootNames[i,1] - BootNames[i,0]
        print(BootNames[i,2])
        print(BootNames[i,2].shape)
        print("done!")
```

In [ ]:

```
#Next we get a mean and standard deviation of each replicate of the pairs, writing it to the SumBootPair Array.

for i in range(NumberPairs):
    BootNames[i,3] = np.zeros([2,Sample])
    TempLen = int(LenArr[i,2])
    TempDF = int(LenArr[i,3])
    for x in range(Sample):
        y = int(x)
        BootNames[i,3][0,y] = np.mean(BootNames[i,2][y,:TempLen])
        BootNames[i,3][1,y] = np.std(BootNames[i,2][y,:TempLen])
    print("Some means from this pair: ")
    print(BootNames[i,3][0,:10])
    print("Some standard deviations from this pair: ")
    print(BootNames[i,3][1,:10])
```

In [ ]:

```
#Almost done! Now we get the means of the means and standard deviations for each pair, sort the means to get a 
#95% confidence interval, and write it all to the Stats array, along with the paired group names.

FinalOutput = np.empty([(NumberPairs + 1), 7], dtype=object)
FinalOutput[0,0] = "Paired Groups"
FinalOutput[0,1] = "Mean of paired difference"
FinalOutput[0,2] = "Mean of standard deviation"
FinalOutput[0,3] = "Control N"
FinalOutput[0,4] = "Treated N"
FinalOutput[0,5] = "CI Max"
FinalOutput[0,6] = "CI Min"

for i in range(NumberPairs):
    row = i + 1
    FinalOutput[row,0] = (DataNames[i, 0] + "_VS_" + DataNames[i,1])
    FinalOutput[row,1] = np.mean(BootNames[i,3][0,:])
    FinalOutput[row,2] = np.mean(BootNames[i,3][1,:])
    FinalOutput[row,3] = LenArr[i,0]
    FinalOutput[row,4] = LenArr[i,1]
    TempSort = np.sort(BootNames[i,3][0,:])
    FinalOutput[row,5] = TempSort[9749]
    FinalOutput[row,6] = TempSort[249]
    
print(FinalOutput)
```

In [ ]:

```
#Last but not least, we will save the final output into a CSV file.

np.savetxt("BootstrapOutput.csv", FinalOutput, delimiter=",", fmt='%s')
```

In [ ]:

In [ ]:

In [ ]:
